# The diversity and ecological significance of microbial traits potentially involved in B_12_ biosynthesis in the global ocean

**DOI:** 10.1101/2023.08.19.553841

**Authors:** Jiayin Zhou, Wei Qin, Xinda Lu, Yunfeng Yang, David Stahl, Nianzhi Jiao, Jizhong Zhou, Jihua Liu, Qichao Tu

## Abstract

Cobalamin (B_12_), an essential nutrient and growth cofactor for many living organisms on the Earth, can be fully synthesized only by selected prokaryotes in nature. Therefore, microbial communities related with B_12_ biosynthesis could serve as an example subsystem to disentangle the underlying ecological mechanisms balancing the function and taxonomy of complex functional assemblages. By anchoring microbial traits potentially involved in B_12_ biosynthesis, we depict the biogeographic patterns of B_12_ biosynthesis genes and their carrying taxa in the global ocean, in light of the limitation to detect *de novo* B_12_ synthesizers via metagenomes alone. Both the taxonomic and functional composition of B_12_ biosynthesis genes were strongly shaped by depth, differentiating epipelagic from mesopelagic zones. The functional genes were relatively stably distributed across different oceans, but their carrying taxa varied considerably, showing clear functional redundancy of microbial systems. Microbial taxa carrying B_12_ biosynthesis genes in the surface water were influenced by environmental factors such as temperature, oxygen and nitrate. However, the composition of functional genes was weakly associated with these environmental factors. Null model analyses demonstrated that determinism governed the compositional variation of B_12_ biosynthesis genes, whereas a higher degree of stochasticity was associated with taxonomic variations. Significant associations were observed between chlorophyll *a* concentration and B_12_ biosynthesis traits, indicating their importance in global ocean primary production. Conclusively, this study revealed an essential ecological mechanism governing the assembly of microbes in nature: the environment selects function rather than taxonomy; functional redundancy underlies stochastic community assembly.

**Impact Statement:** A central question in ecology is how a galaxy of microbial taxa is assembled and distributed across space and through time, executing essential ecosystem functions. By anchoring microbial functional traits potentially involved in B_12_ biosynthesis and their carrying microbial taxa in the global ocean, this study addresses essential ecological questions from functional and taxonomic angles. Integrating multiple lines of evidence, we show that the ecosystem selects functional traits rather than taxonomic groups, and functional redundancy underlies stochastic taxonomic community assembly. Also, microbial communities potentially involved in B_12_ biosynthesis are significantly associated with chlorophyll *a* concentration, demonstrating their importance in global ocean primary production. This study provides valuable mechanistic insights into the complex microbial community assembly in natural ecosystems.

## Introduction

As the home to a galaxy of life forms (371), the global ocean provides roughly 97% of the water on the Earth and 50% of the oxygen, and plays an irreplaceable role in impacting the global climate (2, 3). Microbial communities, the unseen majority (4), are of fundamental importance in maintaining the functionality and stability of the global ocean ecosystems. Not only do they drive the global biogeochemical cycling of various nutrients and elements and maintain ecosystem multi-functioning (5, 6), but also they provide essential nutrients to other organisms, including both prokaryotes and eukaryotes (7). One such example is cobalamin (B_12_), an essential nutrient and growth cofactor that is utilized extensively by prokaryotes and eukaryotes for numerous metabolic functions (8–11). In natural ecosystems, B_12_ biosynthesis is energetically extremely expensive, which causes a high metabolic burden for B_12_ producers (12). Only a small cohort of prokaryotes holds the genetic potential to accomplish such a complex process, while the others have to rely on exogenous supply, forming the “corrinoid economy” (13). Therefore, B_12_ auxotroph may establish close mutualistic interactions with B_12_ producers, offsetting the cost of B_12_ biosynthesis to ensure sustainable sources (14). Such interactive relationships have significant impacts on the composition and structure of marine microbial communities. Two distinct pools of B_12_ analogs were found in the ocean, including the cobalamin pool produced by a few prokaryotes such as Thaumarchaeota and alpha-/gamma-proteobacterial lineages (e.g. *Rhodobacterales*, *Rhizobiales* and most of *Rickettsiales*) (7, 11, 14, 15), and the pseudocobalamin pool produced by Cyanobacteria as representatives (11, 14). Over the past years, the importance of B_12_ has been widely noticed, including influencing the growth rate of phytoplankton in the ocean (16), impacting the size and diversity of microbial community in terrestrial ecosystems (17), and altering the health status of hosts in the human intestinal system (18, 19). In addition, the availability of B_12_ has critical impacts on both the cellular-level metabolic processes (e.g. methionine synthesis) (20), and the system-level biogeochemical cycling (e.g. photosynthesis, aerobic nitrogen cycle) (7, 21, 22). As one of the highly limited nutrients and growth factors controlled by the minority, B_12_ can be considered as a “hard currency” in the global ocean ecosystem.

Over the past years, several studies focusing on the importance of marine B_12_ biosynthesis have been carried out. For example, most of the eukaryotic phytoplanktons in the surface ocean are B_12_ auxotrophic (9), and the growth rate can be limited by the availability of B_12_, further affecting primary productivity. In addition, C:P ratios of B_12_-limited cells in diatom stoichiometry of the Subarctic Pacific are significantly lower in comparison with B_12_-replete cells (23). This phenomenon becomes more pronounced with the significantly increased partial pressure of CO_2_ caused by anthropogenic activities and global climate changes. For example, the C:P ratio gap between B_12_-replete and B_12_-limited cells gradually widens with the increase of carbon dioxide partial pressure (pCO_2_), reaching about 40% at 670 p.p.m pCO_2_ (23). Recent studies have also demonstrated that the growth rate and primary productivity of phytoplankton are affected by the availability of B_12_ (22–24). Although of critical importance, the diversity, distribution and underlying ecological mechanisms shaping the patterns of microbial communities involved in B_12_ biosynthesis in the global ocean remain largely unexplored. This will not only provide valuable insights into a clearer understanding of this subset of prokaryotes in the global ocean, but also shed light on the consequential global ocean ecosystem function. Importantly, the *TARA* Oceans expedition (25–27) provides a valuable resource that includes comprehensive data sets at the global scale, covering a total of eight ocean regions and three ocean depth ranges, making it possible to investigate the global patterns of various microbial (sub)communities, including the microbial taxa related with B_12_ biosynthesis.

In this study, by utilizing the *TARA* Oceans shotgun metagenome sequencing datasets, we surveyed the diversity patterns and ecological importance of microbial traits (functional genes and the corresponding taxonomic groups) potentially involved in B_12_ biosynthesis in the global ocean ecosystem. Community level investigations were mainly performed, owing to the limitations of identifying *de novo* B_12_ synthesizers via metagenomes alone. The following essential ecological questions were addressed: (i) How are B_12_ biosynthesis traits distributed globally? (ii) What ecological mechanism drives and maintains the diversity patterns of B_12_ biosynthesis traits? (iii) How do microbial B_12_ biosynthesis traits contribute to the global ocean ecosystem function, e.g. the ocean’s primary production? Owing to their criticality to the global ocean, we expected that the relative abundance of microbial functional genes involved in B_12_ biosynthesis should be relatively stably distributed in the global ocean. However, owing to the functional redundancy of microbial systems (28), the microbial taxonomic groups carrying them may vary across different oceanic regions and depths. Determinism, therefore, should be mainly responsible for the diversity patterns of functional traits. However, compared to functional traits, microbial taxonomic groups would be relatively more influenced by stochastic processes, due to functional redundancy in microbial systems. Our results well supported the above hypothesis and showed that B_12_ biosynthesis traits are significantly associated with chlorophyll *a* concentration, confirming its important role in ocean primary production.

## Results

### Overall diversity of potential B_12_ biosynthesis traits in the global ocean

B_12_ can be fully synthesized only by a small fraction of prokaryotes (7, 15) due to the multiple enzymatic steps involved (Supplementary Figure 1). By applying VB_12_Path (29) to the *TARA* Oceans shotgun metagenome data, an average of 0.2% reads per sample were identified to encode gene families potentially involved in B_12_ biosynthesis pathways. Consistent with the *TARA* Oceans study that the whole microbial communities significantly differ between mesopelagic layers (MES) and epipelagic zones (26), the same pattern was observed for microbial taxa carrying B_12_ biosynthesis genes. Compared with that in epipelagic zones, the microbial communities potentially involved in B_12_ biosynthesis in MES showed significantly higher taxonomic and functional diversity as well as dramatically different composition (Fig. 1A, Supplementary Figure 2, 3 and 4, Supplementary Table 1). Surprisingly, the evenness of B_12_ biosynthesis functional traits and their carrying taxa were negatively correlated, leading to negatively correlated community diversity (Shannon-Wiener index) (Supplementary Figure 5). The negative correlation is likely due to the fact that only a small fraction of microbial taxa carries a (nearly) full set of gene families involved in B_12_ biosynthesis, thereby even distribution of microbial taxa resulted in uneven distribution of functional traits.

**Figure 1.**
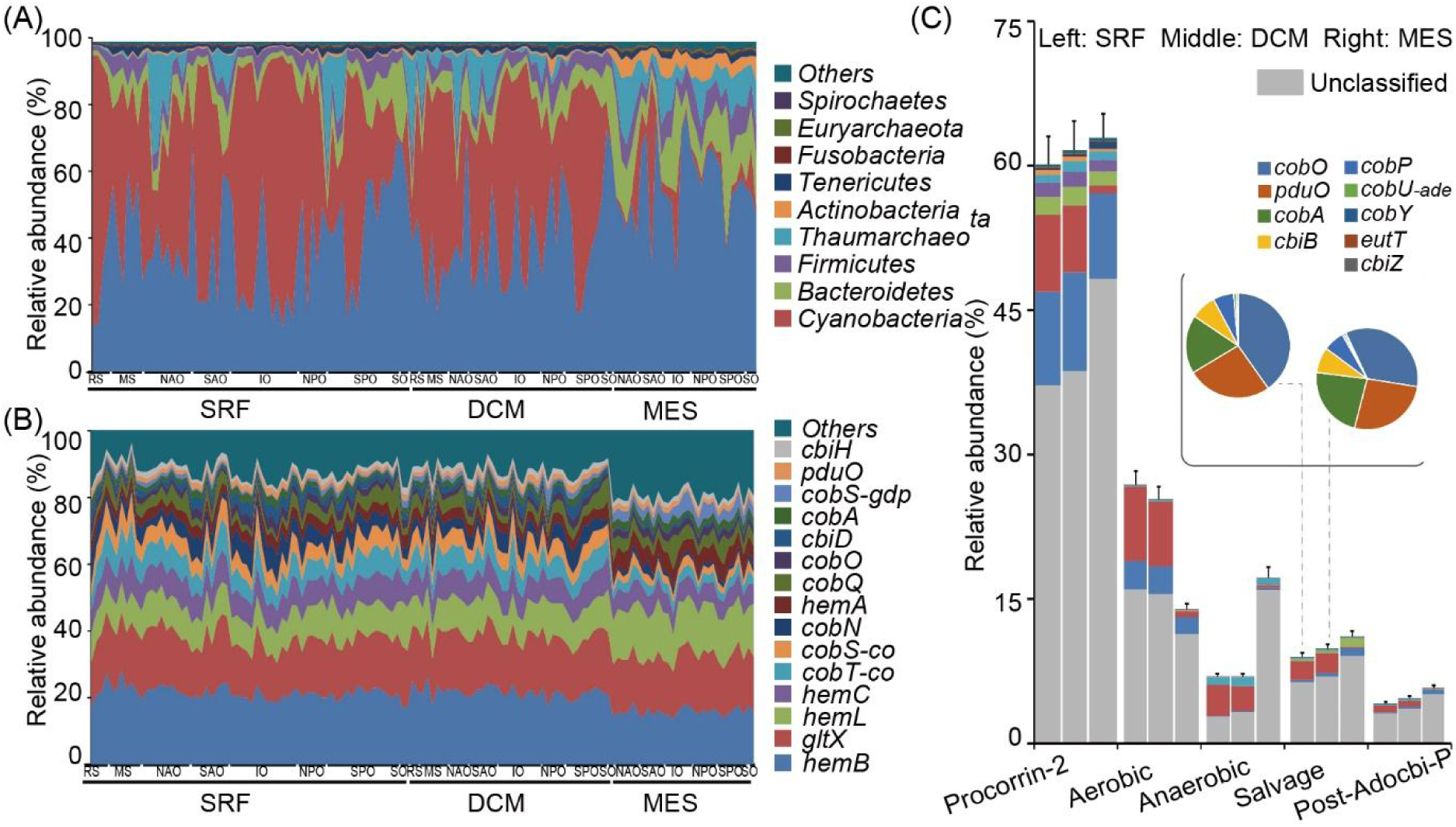
The compositions of microbial taxonomic groups and functional traits related with B_12_ biosynthesis in the global ocean. (A) Compositions of microbial taxa carrying B_12_ biosynthesis genes across different samples; (B) Compositions of microbial functional traits potentially involved in B_12_ biosynthesis across different samples; (C) Relative abundances of microbial phyla carrying genes in different B_12_ biosynthesis pathways and different ocean layers. Relative abundance of functional traits belonging to the salvage pathway in epipelagic zones were presented in pie charts. For the panel A and bar chart in panel C the same scaling color code was used. For the panel B and pie chart in panel C the same scaling color code was used. The major microbial taxa and functional traits were displayed in the figure. SRF, surface water layer; DCM, deep chlorophyll maximum layer; MES, mesopelagic zone.

At the pathway level, microbial functional traits potentially involved in precorrin-2 synthesis (63.84%) and aerobic (24.48%) pathways exhibited the highest relative abundance in the TARA Oceans samples, while anaerobic (9.26%) and post-Adocbi-P (4.87%) pathways were less abundant (Fig. 1C). At the functional gene level, gene families related with aerobic pathways were generally more abundant in epipelagic zones, while the ones related with anaerobic pathways were more abundant in the MES (Fig. 1C, Supplementary Table 4). Most importantly, consistent with our expectations, the relative abundance of functional genes was relatively stable in the global ocean (Fig. 1B), while the taxonomic composition is highly variable. This pattern was observed for microbial communities sampled from different depth intervals and oceanic regions (Fig. 1A). Such results pinpointed an essential microbial ecological discipline that taxonomically highly varied microbial communities still executed similar ecosystem functions.

### Microbial taxa carrying B_12_ biosynthesis genes in the global ocean

Among the identified microbial taxa containing B_12_ biosynthesis genes, Proteobacteria were abundantly detected in all samples, whereas Cyanobacteria dominated in epipelagic zones and dramatically depleted in mesopelagic waters. Compared to that in epipelagic zones, Thaumarchaeota were significantly enriched in the MES, especially targeting the anaerobic pathway of B_12_ biosynthesis, for which 9 *cbi* genes were detected (Fig. 1A and C, Supplementary Table 3). Different modules of B_12_ biosynthesis pathway were featured by different microbial taxonomic groups (Fig. 1C). This was especially evident for samples in the MES. Microbial taxa belonging to Thaumarchaeaota and Bacteroidetes were respectively dominantly observed with genes belonging to anaerobic and salvage pathways. This agrees with previous studies suggesting that cobalamin in the surface ocean may be primarily the result of *de novo* synthesis by heterotrophic bacteria or via modification of cyanobacterially-produced pseudocobalamin, while Thaumarchaeota may be a major cobalamin producer at depth (14). Despite the high abundance of Bacteroidetes in the MES, studies have shown that only 0.6% of Bacteroidetes have complete B_12_ synthesis pathways (15). Gene families (e.g. *cobO*, *pduO* and *cobA*) belonging to the salvage pathway were dominantly carried by Cyanobacteria, more specifically *Prochlorococcus* (Fig. 1C, Supplementary Table 2). A quick BLAST searching these gene families against *Prochlorococcus* genomes in the NCBI database also suggested the wide spreading of these gene families in *Prochlorococcus* (data not shown). While Cyanobacteria are in general pseudocobalamin synthesizers (30), the carrying of gene families belonging to the salvage pathway by *Prochlorococcus* indicated the potential of this genus to remodel B_12_ precursors/analogs in certain conditions. Notably, a recent genomic study also observed salvage pathway gene families in *Synechococcus* genomes, possibly due to horizontal gene transfer events or loss of function (*de novo* B_12_ biosynthesis) during evolution (31). In addition, a high portion of microbial taxa carrying B_12_ biosynthesis genes belonged to unclassified taxonomic groups, especially in the MES, suggesting that much remains to be further explored for the B_12_ biosynthesis gene and taxa in the deep ocean.

Microbial taxa potentially involved in B_12_ biosynthesis in the global ocean were further investigated (Supplementary Table 2) by selecting the putative key B_12_ synthesis gene families in previous investigations (7, 32). B_12_ biosynthesis genes were detected in many microbial taxa, but microbial taxa carrying complete *de novo* B_12_ biosynthesis pathways were rarely found, possibly due to inadequate sequencing depth for detecting these genes and the rarity of microbial taxa containing complete B_12_ biosynthesis pathways. Overall, microbial species including *Prochlorococcus marinus, Candidatus Nitrosopelagicus brevis*, *Candidatus Nitrosomarinus catalina* and *Synechococcus sp. CC9902* were the taxa carrying a high number of key B_12_ biosynthesis gene families. Although B_12_ biosynthesis genes were detected in some microbial taxa (e.g. *Synechococcaceae*, *Prochlorococcaceae* and *Pelagibacteraceae*), they were considered as auxotrophic due to the lack of gene families necessary for DMB such as *bluB* (necessary for DMB biosynthesis) (33, 34) and *cobT* (required for DMB activation) (32). For example, the genus *Synechococcus* contains many genes belonging to B_12_ biosynthesis pathways but lacks key genes for DMB synthesis (Supplementary Table 2) and has been shown to be B_12_ auxotrophic by previous studies (32). Therefore, detection of B_12_ biosynthesis genes in microbial taxa does not necessarily imply the capacity of *de novo* biosynthesis of this cofactor. Further experimental evidence is required to validate such capacity. The results also demonstrated great challenges in identifying potential B_12_ synthesizers using metagenomic approaches, on the basis that the majority of microbial taxa were unknown and metagenomic recovery of rare microbial taxa was almost not possible.

### Latitudinal diversity patterns and distance-decay relationships

We also investigated whether microbial communities potentially involved in B_12_ biosynthesis followed typical biogeographic patterns such as latitudinal diversity gradient (LDG) and distance-decay relationships (DDR), which are well recognized ecological patterns for both microbial and macrobial communities (35, 36). Discordant patterns between the compositions of microbial taxonomic groups and functional genes were observed in this study (Fig. 1A and B). B_12_ biosynthesis serves as an essential ecosystem function and shall be stably maintained in the global ocean. However, the microbial taxa carrying these functional traits are influenced by various environmental conditions. We expected clear LDG and DDR patterns for microbial taxa carrying B_12_ biosynthesis genes, but weaker or even nonexistent patterns for the functional genes. Consistent with our expectation, LDG pattern was weakly observed for the functional genes at the surface water layer (SRF) (*P* = 0.02), and not found at the deep chlorophyll maximum layer (DCM) and mesopelagic zone (MES). No significant DDR pattern was observed for B_12_ biosynthesis genes at all three pelagic zones. For microbial taxa carrying B_12_ biosynthesis genes, a strong LDG pattern was observed at the SRF zone (*P* = 0.006). DDR, however, was observed at all three pelagic zones (*P* ≤ 1e-7) (Fig. 2AB). Such distinct biogeographic patterns of functional genes and taxonomic groups again pointed to the essential microbial ecology principle, i.e. microbial functional genes executing essential ecosystem functions are prevalently distributed, whereas their carrying microbial taxa may dramatically vary.

**Figure 2.**
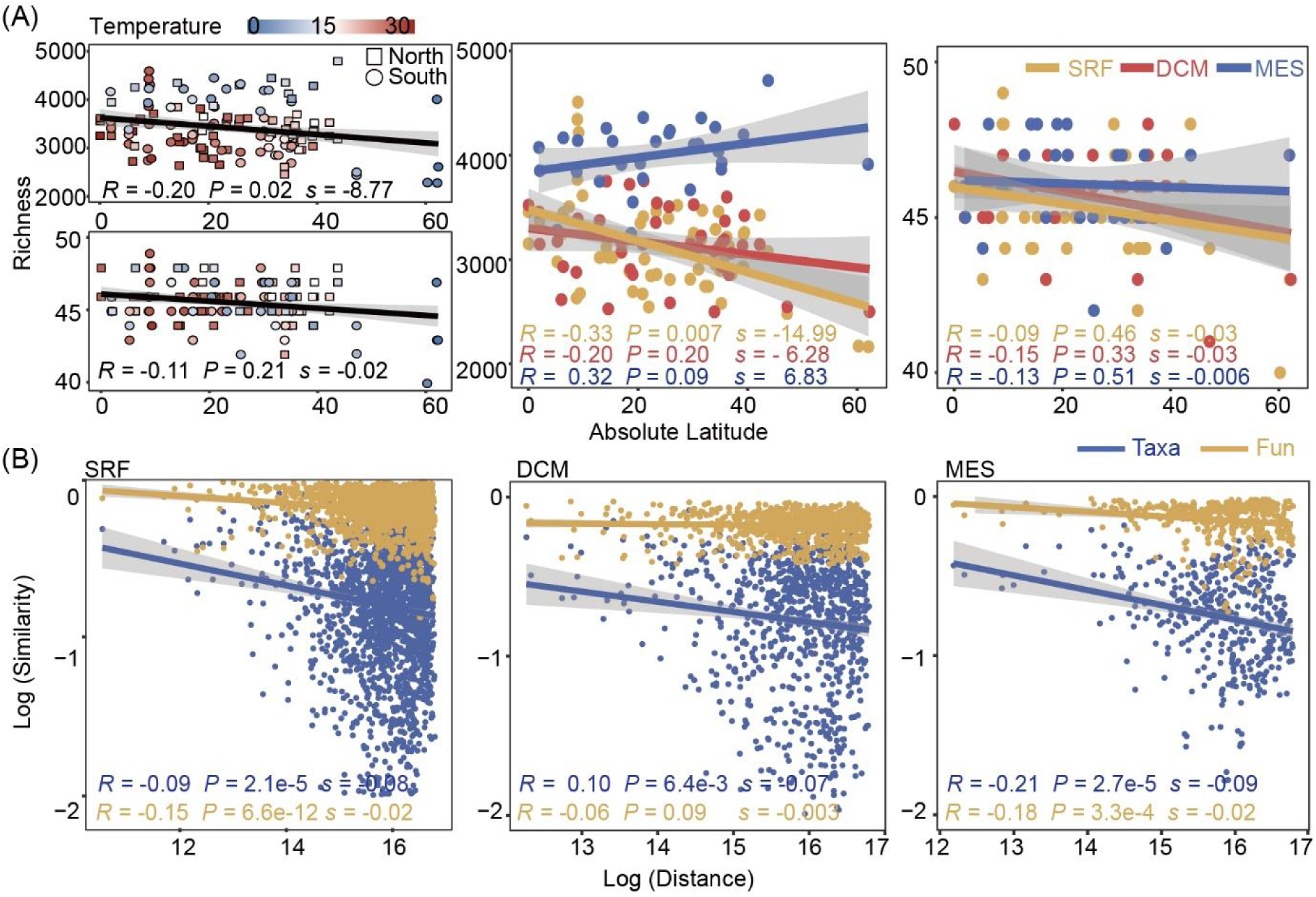
Biogeographic patterns of potential B_12_ biosynthesis traits in the global ocean. (A) Latitudinal diversity gradient patterns for B_12_ biosynthesis traits in the global ocean; (B) Distance-decay relationship for B_12_ biosynthesis traits in the global ocean. The patterns for taxonomic groups and functional traits were investigated. Taxa, taxonomic composition; Fun, functional composition.

### Environmental factors associated with the variations of potential B_12_ biosynthesis traits

Here, the associations between B_12_ biosynthesis traits and environmental factors were also investigated (Supplementary Figure 6). Since both the functional and taxonomic compositions of B_12_ biosynthesis genes dramatically differ by depth, the associations with geo-environmental factors were analyzed by a given range of water depths, eliminating the effects of depth and depth-correlated environmental factors. As a result, weakened effects of environmental factors on the taxonomic compositions were observed from the SRF to the MES layers. In the SRF layer, the concentrations of dissolved oxygen and nitrate availability were significantly associated with the taxonomic compositions. Such effects, however, gradually diminished in the DCM and MES layers. Interestingly, significant associations were not observed between environmental factors and the functional compositions of B_12_ biosynthesis genes in all three oceanic layers, suggesting that changes in environmental conditions mainly affected the taxonomic compositions.

The associations between environmental factors and community diversity were also investigated. Significant associations between environmental factors and community diversity could be observed (Supplementary Figure 7A). However, such effects were weakened or even diminished when looking at individual pelagic zones (Supplementary Figure 7B, C, and D), suggesting that depth differences from SRF to MES layers and their correlation with environmental factors were mainly responsible for such “pseudo-associations”. Surprisingly, the effects of temperature on B_12_ biosynthesis functional trait diversity differed dramatically by oceanic layers. The temperature was positively associated with the functional gene diversity in epipelagic layers (Supplementary Figure 7B and C), but negatively in the MES (Supplementary Figure 7D), leading to insignificant associations across the whole upper ocean (Supplementary Figure 7A). Such opposite patterns were also observed for other environmental factors such as oxygen, nitrite and nitrate concentration (NO_2_NO_3_), and nitrate, though some of them were not statistically significant (*P* ≥ 0.05).

### Ecological mechanisms governing the assembly of B_12_ biosynthesis traits

Considering the critical roles that B_12_ play in the ecosystem, we expected that the assembly of microbial functional traits be highly deterministic. To examine this hypothesis, we quantified the relative importance of deterministic and stochastic processes in governing the assembly of functional traits potentially involved in B_12_ biosynthesis. Here, the null model analysis was employed to characterize the ratio of stochasticity to determinism by comparing the observed and null model community β-diversity (Fig. 3A). Consistent with our hypothetical expectations, the stochastic ratio suggested that both the assembly of microbial functional genes and their carrying taxa were highly deterministic. Compared to the functional traits, the taxonomic groups had higher stochastic ratios, especially in the MES layer, suggesting that the assembly of taxonomic groups was more stochastic than functional traits. Such patterns of stochastic ratios between functional traits and taxonomic groups were consistent in different oceanic layers.

**Figure 3.**
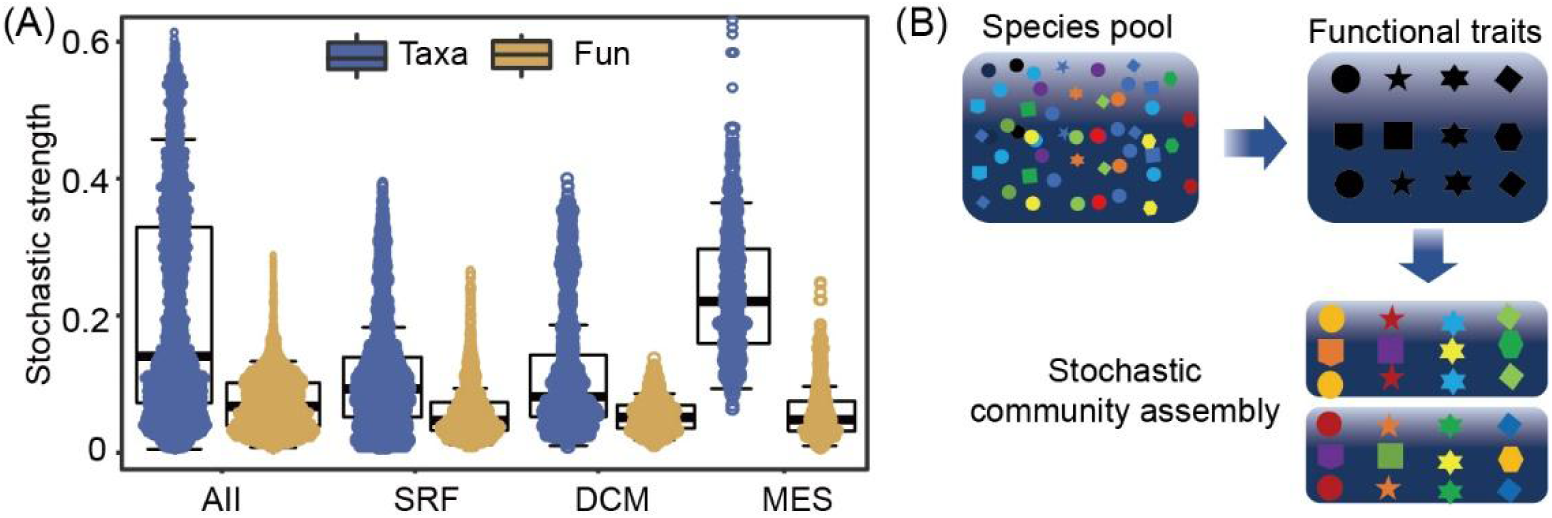
Mechanisms governing the assembly of B_12_ biosynthesis traits in the ocean ecosystem. (A) Stochasticity of community assembly as revealed by null model analysis; (B) An ecological model explaining community assembly of microbial functional groups in the ocean ecosystem. According to the model, the environment selected microbial functional traits rather than taxonomy, and functional redundancy underlies stochastic community assembly. In the ecological model, different colors represent different microbial taxa, whereas different shapes represent different functional traits. Taxa, taxonomic composition; Fun, functional composition.

We hypothesized that deterministic factors should govern the assembly of microbial functional traits, and that the assembly of microbial taxa shall be relatively more stochastic than functional traits. All the results described above, e.g., stable distribution of functional traits vs. highly varied taxonomic groups (Fig. 1AB), stronger biogeographic patterns for taxonomic groups than functional traits (Supplementary Figure 6 and Supplementary Figure 7), and relative importance of deterministic and stochastic processes (Fig. 3A), provided supportive evidence for our hypotheses for community assembly of B_12_ biosynthesis traits. Integrating all lines of evidence, we proposed a functional-trait-based ecological model to explain the complex microbial community assembly in natural ecosystems (Fig. 3B). Variations of geo-environmental factors such as depth, temperature, and oxygen form multiple ecological niches in the oceanic ecosystem (e.g. epipelagic zone and MES). Microorganisms capable of living in these ecological niches comprise the species pools. To maintain fundamental ecosystem function, microorganisms carrying essential functional traits are selected. Therefore, it is the function rather than taxonomy that the environment truly selects (37). However, owing to functional redundancy in microbial systems (28), different taxonomic groups carry the same functional traits. Meanwhile, stochastic processes such as drift and dispersal are associated with microbial taxa. Stochastic community assembly occurs at the time these functional traits are selected. As a result, varied taxonomic compositions come with comparable functional traits combinations, as have been observed in multiple ecosystems (38–40). For microbial traits potentially involved in B_12_ biosynthesis, both taxonomic groups and functional traits were governed by deterministic processes, and functional redundancy of microbial taxonomic groups led to higher stochasticity in community assembly.

### Ecological importance of potential B_12_ biosynthesis traits in the global ocean

Finally, we investigated the ecological roles that the potential B_12_ biosynthesis traits play in the oceanic ecosystem, such as cobalamin-dependent microorganisms and their contribution to the ocean’s primary production (7, 9, 14, 24). To investigate if B_12_ biosynthesis traits are potentially associated with cobalamin-dependent microbial communities and global ocean primary productivity, we investigated the associations between the community diversity of B_12_ biosynthesis traits with the relative abundances of the *metH* gene family (encoding cobalamin-dependent methionine synthase) and the chlorophyll *a* concentration. First, a significant association was observed between the relative abundance of *metH* gene family and B_12_ biosynthesis trait diversity (Supplementary Figure 8), confirming the importance of B_12_ biosynthesis members to cobalamin-dependent members in the oceanic ecosystem. Second, the concentrations of chlorophyll *a* in the epipelagic zone were also significantly associated with B_12_ biosynthesis trait diversity (*P* ≤ 0.005) (Fig. 4A). Notably, the concentrations of chlorophyll *a* were positively correlated with the taxonomic diversity of B_12_ biosynthesis traits, but negatively with functional gene diversity. Such an opposite pattern was attributed to the negative correlation between the evenness of B_12_ biosynthesis genes and their carrying taxa (Supplementary Figure 8). Meanwhile, to exclude the potential influence of the whole microbial community and further confirm the significant correlations between chlorophyll *a* concentration and B_12_ biosynthesis traits, we also inspected the associations between the concentrations of chlorophyll *a* and the prokaryotic community diversity. As a result, the association strength between chlorophyll *a* concentrations and prokaryotic community diversity was either insignificant or much weaker than that with B_12_ biosynthesis traits (Fig. 4B). Finally, the machine learning approach random forest was employed to further verify the importance of B_12_ biosynthesis traits by predicting chlorophyll *a* concentration using B_12_ community profiles. The results demonstrated that both the taxonomic and functional profile of B_12_ biosynthesis traits can well predict the concentration of chlorophyll *a* in the ocean (Fig. 4C and D). This also held true when using SRF microbial data as training data set and predicting chlorophyll *a* in the DCM layer, or vice versa (Supplementary Figure 9).

**Figure 4.**
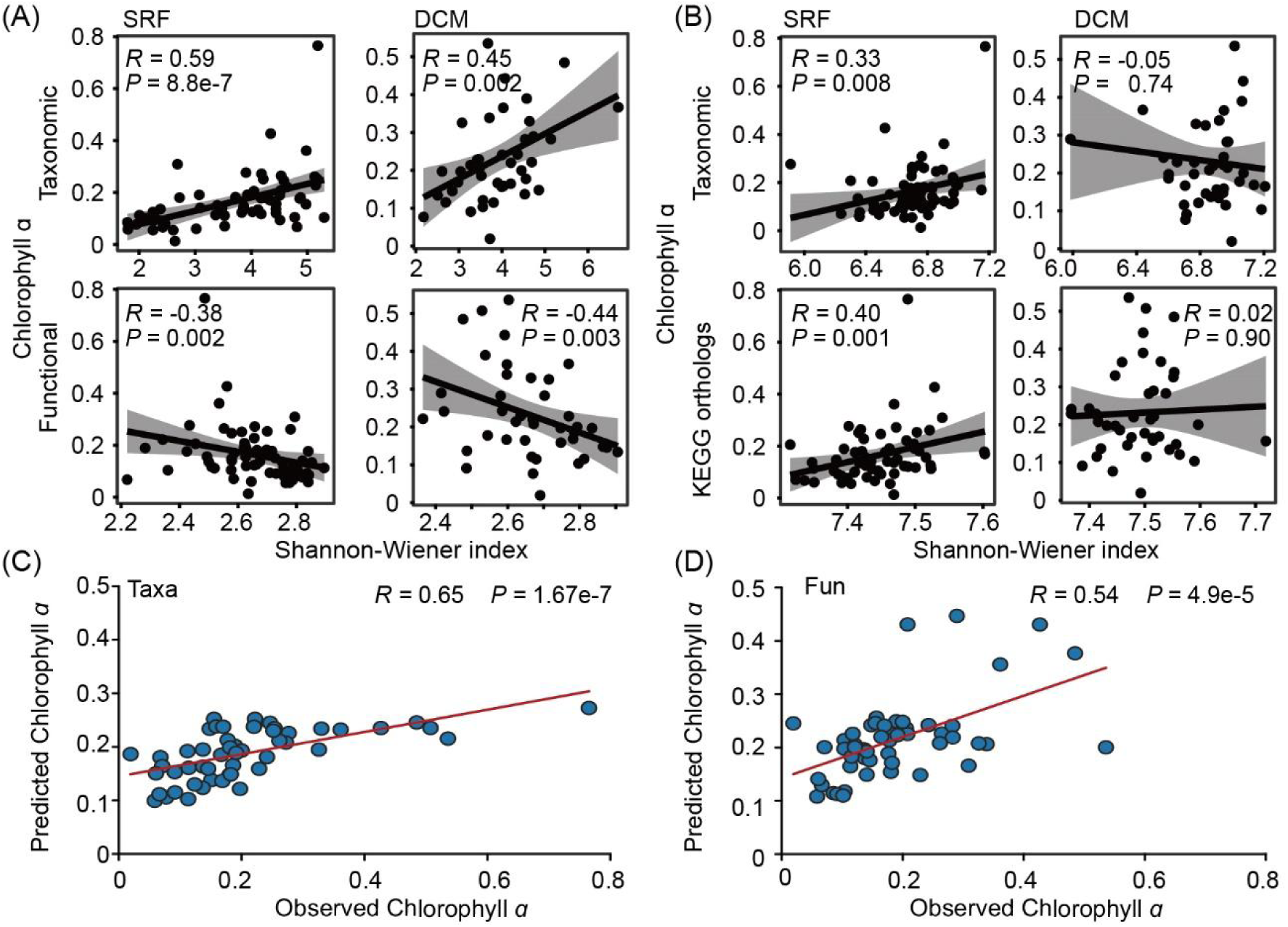
Association between microbial community diversity and chlorophyll *a* concentrations in the global ocean. (A) Associations (Spearman’s ρ) between B_12_ biosynthesis trait diversity (taxonomic and functional trait) and chlorophyll *a* concentrations; (B) Associations (Spearman’s *ρ*) between overall prokaryotic community diversity (taxonomic and KEGG orthologous groups) and chlorophyll *a* concentrations; (C) Predicted chlorophyll *a* concentrations using microbial taxa carrying B_12_ biosynthesis genes; (D) Predicted chlorophyll *a* concentrations using B_12_ biosynthesis functional trait profiles. Taxa, taxonomic composition; Fun, functional composition.

## Discussion

Focusing on “who is doing what, where, and how?”, this study investigated the ecological mechanisms driving the diversity patterns of microbial traits potentially involved in B_12_ biosynthesis and their ecological importance in the global ocean. Limited by the rarity of the targeted microbial taxa and current technologies, confident inference of *de novo* B_12_ synthesizers can hardly be approached. Therefore, community-level investigations were performed in this study. Similar to what has been observed for the global ocean microbiome (26), both the taxonomic and functional gene composition related with B_12_ biosynthesis differed by depth instead of oceanic regions. Multiple factors such as depth, light and temperature and other associated environmental factors shall be responsible for such patterns. This probably suggested completely different niche preferences by B_12_ biosynthesis traits in different oceanic layers. We also noticed that the evenness of B_12_ biosynthesis genes and their carrying taxa were negatively correlated, suggesting that even distribution of microbial taxa may not lead to even distribution of functional traits. Such negative correlation should be due to the fact that only a small fraction of microbial taxa contains (near) complete B_12_ biosynthesis pathway in their genomes, and even distribution of microbial taxa does not reflect even functional traits.

Microbial taxa carrying B_12_ biosynthesis genes in the ocean ecosystem were also investigated at a refined taxonomic resolution. However, limited information could be gained. First, the taxonomy of the majority of B_12_ biosynthesis genes remained unclassified, even against taxonomic databases built from the most recent NCBI database. This is especially crucial for microbial taxa in the MES. Such shortage is mainly caused by the limitations of current genomic databases (41), the uncultured majority of microbial taxa in nature (42), and potential limitations of read-based analyses. This also suggests that much remains to be further explored for this tiny group of microorganisms on Earth, especially in the deep ocean. Second, consistent with our current knowledge (14), only a few microbial genera in the ocean were found to carry *de novo* B_12_ biosynthesis potential by judging the gene families linked to the microbial taxa. However, comparative genomics of currently sequenced microbial genomes from NCBI RefSeq suggest that 37% of prokaryotic microbial species have the potential to biosynthesize cobamides *de novo*, though complete pathways are not always detected (15). Among these, 57% of Actinobacteria are predicted to biosynthesize cobamides, whereas only 0.6% of Bacteroidetes have the complete pathway (15). Such inconsistency between metagenomic and genomic studies indicated the rarity and unknown property of *de novo* B_12_ synthesizers in the ocean, and current sequencing technologies and depth may not well capture them. Third, identifying *de novo* B_12_ synthesizers is challenging and requires further attention. *Rhodobacteraceae*, *Rhizobiales* and a subset of Cyanobacteria were the most important candidates to be B_12_ prototrophic in neritic ecosystems by metatranscriptomic and metaproteomic analyses (43). However, one needs to be aware that the lower ligand is DMB to produce B_12_ and not pseudocobalamin. Perhaps judgment by synthesis and activation of key genes of DMB, e.g., *bluB* (33, 34) and *cobT* (32), is also an option. Cyanobacteria strains release pseudo-B_12_ into the media at a high rate, and it is speculated that Cyanobacteria may be the main providers of (pseudo-)B_12_ in algae metabolism (44). Similarly, genes potentially involved in B_12_ biosynthesis were frequently detected in Cyanobacteria genera such as *Synechococcus* and *Prochlorococcus*, which may only produce pseudocobalamin due to adenine as the low ligand instead of DMB, consistent with previous studies (7, 11, 14). In certain cases, microbial taxa (e.g. *Dehalococcoides mccartyi strain 195*, *Chlamydomonas reinhardtii*) may remodel non-functional cobamides (e.g. pseudocobalamin) to B_12_ under suitable environmental conditions such as at the presence of DMB or its intermediate α-ribazole (11, 32, 45). Interestingly, *bluB* and *cobT* were detected from *Prochlorococcus marinus* at high taxonomic levels (Supplementary Table 2), and previous studies also mentioned that *Prochlorococcus marinus SS120* may encode the full set of enzymes of the biosynthetic pathways for heme B_12_ (46). In the marine ecosystem, *Rhodobacterales* are the major alphaproteobacterial B_12_ producers, but we did not detect *bluB* from it (e.g. *Epibacterium mobile*). Therefore, even if these key B_12_ biosynthesis gene families are detected, further experimental validation is needed for confirming their function in the ecosystem.

This study also revealed important implications regarding the ecological roles that B_12_ biosynthesis traits play in the oceanic ecosystem. Previous studies suggested that eukaryotic phytoplankton in the surface ocean are B_12_ auxotrophs (9, 32), and the growth rate may be limited by the availability of B_12_, further affecting ocean primary productivity (16, 24, 47, 48). Most of these eukaryotic algae requirements for B_12_ are primarily mediated by methionine synthase (9, 49), a key enzyme in cellular one-carbon metabolism responsible for catalyzing the conversion of homocysteine and 5-methyl-tetrahydrofolate to tetrahydrofolate and methionine (50, 51). Although B_12_-independent methionine synthase (MetE) and B_12_-dependent methionine synthase (MetH) are capable of completing this reaction (9, 49), MetE is approximately 100-fold less efficient in catalysis than MetH (52), and this inefficiency further results in an approximately 30-40-fold increase in nitrogen and zinc requirements for MetE compared to MetH (53). Consistent with previous studies, significant correlation was observed between B_12_ biosynthesis traits and *metH* gene encoding B_12_-dependent methionine synthase, as well as chlorophyll *a* concentration. This suggested that B_12_ biosynthesis traits exerted strong effects on chlorophyll *a* concentration, demonstrating the importance of this microbial group on primary production in the global ocean.

In addition to investigating the diversity patterns of B_12_ biosynthesis traits in the oceanic ecosystem, the microbial subcommunities also served as an example to reveal an intriguing functional-trait-based ecological mechanism explaining the complex microbial community assembly in nature. Both deterministic and stochastic processes govern microbial community assembly, and a major question is which one is more important (54–56). Considering that B_12_ biosynthesis is an essential ecosystem function and shall be stably maintained in the global ocean (7, 15, 57), we speculated that strong determinism shall govern the assembly of potential B_12_ biosynthesis traits. However, microbial communities are usually functionally redundant (28), for which multiple different microbial taxa may execute the same function. Similar to previous studies in the ocean (26, 58, 59), high functional redundancy was also observed in this study. A previous study suggests that the ecosystem tends to select microbial functional traits rather than taxonomic groups (37). In addition, stochastic processes such as drift and dispersal are associated with microbial taxa (60). As multiple microbial taxa carried the same functional traits, a certain degree of randomness shall be associated with microbial taxa in the ecosystem. Consistent with our expectations, higher stochasticity was observed in the assembly of microbial taxa than functional traits. To summarize, the environment selects microbial functional traits rather than taxonomy (37), and functional redundancy (28) underlies stochastic microbial community assembly, thereby maintaining essential ecosystem function and stability (61). In addition, we urge mechanistic studies in microbial community ecology should not only focus on microbial taxonomy, but also functional genes that they carry. Whenever possible, microbial functional genes and taxonomy shall be equally considered in microbial systems.

In conclusion, using B_12_ biosynthesis subsystem as an example, this study investigated the diversity, biogeographic patterns, and ecological drivers of this specific microbial functional group in the global ocean. Comparative pattern analyses of B_12_ biosynthesis genes and their carrying microbial taxa revealed an important microbial ecological mechanism, elucidating the relationship between natural ecosystems and complex microbial communities from the functional angle. Also, B_12_ biosynthesis traits were significantly associated with chlorophyll *a* concentration, demonstrating its importance in global ocean primary production. This study provided valuable mechanistic insights into the complex microbial community assembly in natural ecosystems.

## Materials and Methods

### Tara Oceans shotgun metagenomes and geo-environmental factors

A total of 359 shotgun metagenomes targeting 138 samples covering three oceanic layers, including surface water layer (SRF, 5 to 10 m), deep chlorophyll maximum layer (DCM, 17 to 180 m) and mesopelagic zone (MES, 250 to 1000 m), were downloaded from the European Bioinformatics Institute (EBI) repository under project ID ERP001736 (26). Forward and reverse reads were merged into longer sequences by the program PEAR (version 0.9.6, -q 30) (62). An average of 208,881,758 merged reads per sample were obtained. Geo-environmental factors, the overall taxonomical profiles, and KEGG orthologous group profiles associated with the shotgun metagenome data were downloaded from http://ocean-microbiome.embl.de/companion.html. Metadata for chlorophyll *a* concentrations in these TARA Oceans samples were obtained from the ZENODO website under the record number 7739198 (https://zenodo.org/record/7739198) according to a previous study (63).

### Metagenomic profiling of marine functional genes potentially involved in B_12_ biosynthesis

To keep the fidelity of taxonomic and functional profiles and get more usable information from the metagenomic dataset (64), read-based analysis was performed. Considering the accuracy of gene definition and computational efficiency, VB_12_Path (29), a specific functional gene database for metagenomic profiling of gene families involved in B_12_ biosynthesis pathways, was employed. Although this database is relatively small, both targeted gene families and their homologs from large public databases (e.g. KEGG, eggNOG and COG) are integrated, minimizing false positive assignments. Briefly, merged metagenomic reads were searched against VB_12_Path. A total of 54 gene families involved in five modules of B_12_ biosynthesis pathway as previously described (29), including precorrin-2 synthesis processes, aerobic pathway, anaerobic pathway, salvage and remodeling pathway, post-Adocbi-P pathway, are targeted in the database. The program DIAMOND (version 0.9.25, option: -k 1 -e 0.0001) (65) was used to search nucleotide sequences against VB_12_Path using the blastx mode. Sequences matching VB_12_Path were retrieved to generate functional profiles targeting gene families involved in marine B_12_ biosynthesis using the PERL script provided in VB_12_Path. To minimize bias associated with sequence number variations across different samples, rarefaction was applied to each metagenome by a random subsampling effort of 100,000,000 sequences. Four samples were excluded from further analysis due to insufficient sequences.

To obtain taxonomic profiles for microbial taxa carrying B_12_ biosynthesis genes, merged metagenomic sequences belonging to targeted gene families in VB_12_Path were extracted by the seqtk program (https://github.com/lh3/seqtk). Extracted sequences were then subject to taxonomic assignment by Kraken2 (66). A standard Kraken2 database was built locally based on the most recent NCBI database at the time this study was carried out. Taxonomic profiles were generated at multiple taxonomic levels based on the Kraken2 report files. After obtaining the functional and taxonomic profiles, Kruskal-Wallis test was conducted to estimate statistical differences in relative abundances of potential B_12_ biosynthesis taxonomic groups and functional traits between the epipelagic (SRF/DCM) zone and MES. The false discovery rate (FDR) approach was employed to adjust the *P* value to control for false positives using the “stats” package in R. All gene families of the B_12_ biosynthetic pathway, and the microbial taxa containing B_12_ biosynthetic gene families are collectively referred to as B_12_ biosynthesis traits in the context.

### Diversity indices

Various diversity indices were calculated by the “vegan” package (67) in R (software version 4.0.3). Specifically, the richness, Shannon-Wiener index and Pielou’s evenness index were calculated for within sample diversity, i.e. alpha diversity. The Bray-Curtis dissimilarity was calculated to represent between sample diversity, i.e. community dissimilarity or beta diversity. The complement of community dissimilarity (1-dissimilarity) was calculated to quantitate community similarity. Both within sample and between sample diversity indices were calculated for functional and taxonomic profiles. Compositional variance among samples in different layers and oceans, as well as epipelagic zone and MES were calculated using Bray-Curtis dissimilarities and explored by Principal coordinated analysis (PCoA), of which the first two axes were extracted for visualization. Three different nonparametric analyses, including permutational multivariate analysis of variance (PERMANOVA), analysis of similarity (ANOSIM) and multi-response permutation procedure (MRPP), were performed to evaluate the statistical significance of compositional variations among SRF, DCM, and MES layers.

### Latitudinal diversity gradient and distance decay relationship

Two major biogeographic patterns, including the latitudinal diversity gradient (LDG) and distance decay relationships (DDR), were analyzed to investigate the diversity trend of B_12_ biosynthesis traits. For LDG, the relationship between community richness (species and functional traits) and absolute latitude was analyzed. For DDR, the relationship between community similarity and geographic distance was analyzed. The geographic distance between different samples was calculated by the Vincenty Ellipsoid formula based on the latitude and longitude coordinates using the “geosphere” package in R (68). Community similarity values (Bray-Curtis indices) were obtained by subtracting community dissimilarity from 1. For DDR analyses, both the geographic distance and community similarity values were logarithmically transformed. For both LDG and DDR, linear regression analysis was carried out to visualize the diversity trendline. Values including correlation coefficients, slope and significance *P* values were calculated. Analyses were performed for samples in three different layers.

### Correlating environmental factors with the diversity and composition of microbial communities

To identify the potential environmental factors shaping the variations of B_12_ community diversity and composition, the partial mantel test was performed by correcting for geographic distance. Bray-Curtis dissimilarity was selected to characterize the community distance for both taxonomic and functional trait profiles. The Euclidean distance method was used to characterize the distance between environmental factors. A permutation time of 9,999 was set for the partial mantel test. A total of 11 environmental variables were recruited, including latitude, longitude, depth, temperature, oxygen, mean nitrates concentration, NO_2_, nitrite and nitrate concentration (NO_2_NO_3_), phosphate (PO_4_), salinity, and silica (Si). To analyze the associations between environmental factors and community diversity, redundancy analysis (RDA) was used to evaluate the collinearity between environmental variables and the taxonomic and functional trait composition. After excluding variables with high collinearity, a total of six geo-environmental variables were retained, including depth, temperature, oxygen, nitrates, NO_2_NO_3_, and PO_4_. Then, linear regression analyses were conducted to investigate the relationships between each remaining individual environmental variable and community diversity (Shannon-Wiener index). Spearman’s rank coefficient of correlation was calculated. All of the above statistical analyses were performed using the “vegan” package (67) in R.

### Correlating *metH* gene abundance and chlorophyll *a* concentrations with B_12_ biosynthesis trait diversity

To disentangle the potential effects of B_12_ biosynthesis traits on cobalamin-dependent microbial communities and the ocean’s primary productivity, the *metH* gene relative abundance and chlorophyll *a* concentration were correlated with the community diversity of B_12_ biosynthesis traits. Of these, *metH* gene was selected for its encoding of B_12_-dependent methionine synthase, a pivotal enzyme of cellular one-carbon metabolism and DNA synthesis (49). Positive associations were expected between *metH* communities and B_12_ biosynthesis functional genes. And chlorophyll *a* was selected as a proxy for phytoplankton biomass to further approximate primary productivity. Linear regression analysis was used to explore the relationship between *metH* relative abundance, the chlorophyll *a* concentration and B_12_ biosynthesis trait diversity. To eliminate the potential impact of the whole prokaryotic community and confirm the importance of B_12_ biosynthesis traits, linear regression analysis was also carried out between the whole prokaryotic microbial community and chlorophyll *a* concentration. Both the taxonomic profiles and functional profiles (KEGG orthologous groups) were analyzed. The analyses were carried out for samples in different layers. Spearman’s rank coefficient of correlation was calculated. Correlation coefficients with significance *P* values < 0.005 was termed as significant correlation.

In addition to linear regression analyses, the machine learning approach random forest was also employed to verify the importance of B_12_ biosynthesis traits on chlorophyll *a* concentration by predicting chlorophyll *a* concentrations using the functional and taxonomic profiles of B_12_ biosynthesis traits. Here, half of the microbial data from epipelagic zones were randomly selected for developing a random forest training model, which was used to predict chlorophyll *a* concentration using the remaining microbial data in epipelagic zones. In addition, individual layers were also validated, using samples from one layer (SRF/DCM) as the training set to predict the chlorophyll *a* concentration in the other layer. The relationship between predicted and observed chlorophyll *a* concentration was analyzed to evaluate the importance of B_12_ communities. The random forest analysis was performed using the “randomForest” package (69) in R.

### Community assembly mechanisms

The null model analysis was employed to investigate the potential ecological mechanisms governing the compositional variations of B_12_ biosynthesis traits. Since the taxonomic and functional trait profiles for B_12_ biosynthesis genes were obtained by extracting targeted sequences from the shotgun metagenomic dataset, phylogenetic markers for these profiles were not applicable. Therefore, the approach proposed by Zhou et al. was employed in this study (70, 71). In the analysis, stochastic strength was calculated via null models to characterize the relative importance of deterministic and stochastic processes in driving the assembly of B_12_ biosynthesis traits. The within-sample (local) and across-sample (regional) richness were constrained to produce null models, in order to rule out potential influence of local and regional species richness on β-diversity (72). A dissimilarity matrix was calculated based on Bray-Curtis index. The complementary similarity matrix was obtained by (1-dissimilarity). This procedure was repeated 1,000 times to generate a total of 1,000 null models, based on which an average similarity matrix was obtained. Community assembly stochasticity was estimated by comparing the observed and randomized community similarity, according to a modified method as described previously (54, 73). The stochastic ratio was calculated considering two scenarios: i) communities are governed by deterministic factors that produce more similar communities. In such case, the observed community similarity (*C*_*ij*_) between the *i*-th and *j*-th communities would be larger than the null expectations (*E͞*_*ij*_). ii) communities are governed by deterministic factors making communities more dissimilar. As such, *C*_*ij*_ would be smaller than *E͞*_*ij*_. As a result, the observed dissimilarity (*D*_*ij*_ = 1 − *C*_*ij*_) would be larger than the null model dissimilarity (*G͞*_*ij*_ = 1 − *E͞*_*ij*_). Hence, the following functions can be used to evaluate the stochastic ratio:

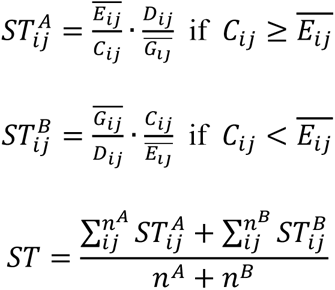

The null model analysis was carried out for both taxonomic and functional profiles. R packages including vegan (67), bioenv (74), and NST (39) were used in the analysis.

## Data availability

Sequences belonging to the B_12_ biosynthesis traits extracted from the TARA Oceans shotgun metagenome datasets are deposited at ZENODO website under the record number 7520550.

## Acknowledgements

This study was supported by National Key Research and Development Program of China (2020YFA0607600, 2019YFA0606700, 2017YFA0604300), the National Natural Science Foundation of China (31971446, 92051110), the Natural Science Foundations of Shandong Province (2020ZLYS04, ZR2020YQ21), the Taishan Young Scholarship of Shandong Province, and the Qilu Young Scholarship of Shandong University. The funders had no role in study design, data collection and interpretation, or the decision to submit the work for publication.

## Author Contributions

Qichao Tu conceived the research. Jiayin Zhou and Qichao Tu performed the bioinformatics analyses and wrote the manuscript. Wei Qin and Xinda Lu helped with the data analysis. Wei Qin, Xinda Lu, Yunfeng Yang, David Stahl, Nianzhi Jiao, Jizhong Zhou and Jihua Liu revised the manuscript.

## Ethics Statement

This study does not contain any studies with human participants or animals performed by any of the authors.

