## Supplementary Figure for "The diversity and ecological significance of microbial traits potentially involved in B_12_ biosynthesis in the global ocean"

**Running title**: $B_{12}$ biosynthesis traits in the global ocean

Jiayin Zhou[1,2], Wei Qin[3], Xinda Lu[4,†], Yunfeng Yang[5], David Stahl[6], Nianzhi Jiao[2,7], Jizhong Zhou[3,8], Jihua Liu[1,2*], Qichao Tu[1,2*]

[1] Institute of Marine Science and Technology, Shandong University, Qingdao, China

[2.] Joint Lab for Ocean Research and Education at Dalhousie University, Shandong University and Xiamen University, Qingdao, China

[3.] Department of Microbiology and Plant Biology, University of Oklahoma, Norman, OK, USA

[4.] Department of Civil and Environmental Engineering, Massachusetts Institute of Technology, Cambridge, MA, USA

[5.] State Key Joint Laboratory of Environment Simulation and Pollution Control, School of Environment, Tsinghua University, Beijing, China

[6.] Department of Civil and Environmental Engineering, University of Washington, Seattle, WA, USA

[7.] Institute of Marine Microbes and Ecospheres, Xiamen University, Xiamen, China

[8.] Earth and Environmental Sciences, Lawrence Berkeley National Laboratory, Berkeley, CA, USA

†. Current affiliation: DermBiont Inc., Boston, MA, USA

### Supplementary Figures

**Supplementary Figure 1.** Cobalamin biosynthesis pathway and related gene families. The $B_{12}$ biosynthesis pathway contains five modules, including precorrine-2 synthesis, aerobic pathway, anaerobic pathway, salvage and remodeling pathway, and post-AdoCbi-P pathway.

**Supplementary Figure 2.** Relative abundances of $B_{12}$ biosynthesis traits in the ocean. The size of the circles and colors represents the relative abundance of microbial functional genes involved in $B_{12}$ biosynthesis.

**Supplementary Figure 3.** Diversity indices (richness, evenness, and Shannon-Wiener index) of microbial communities carrying $B_{12}$ biosynthesis in SRF, DCM, and MES layers. Both the diversity indices for taxonomic groups (top) and functional traits (bottom) were presented.

**Supplementary Figure 4.** PCoA clustering of taxonomic and functional trait potentially involved in $B_{12}$ biosynthesis in the global ocean. Bray-Curtis dissimilarity was used for distance calculation. Different colors represent samples in different ocean layers, while different shapes represent different oceans.

**Supplementary Figure 5.** Relationship between $B_{12}$ biosynthesis functional traits and their carrying taxa in different oceanic layers. Diversity indices including richness, evenness, and Shannon-Wiener index were analyzed.

**Supplementary Figure 6.** Linking environmental factors with the compositional variations of $B_{12}$ biosynthesis traits. Pairwise comparisons of environmental factors are shown, with color gradients denoting Spearman's correlation coefficients. The $B_{12}$ biosynthesis traits in the SRF (A), DCM (B), and MES (C) layers were related to each environmental factor by partial (geographic distance–corrected) Mantel tests. Edge width corresponds to Mantel's r statistic, edge color corresponds to the statistical significance based on 9,999 permutations. Taxa: taxonomic composition; Fun: functional composition.

**Supplementary Figure 7.** Associations (Spearman's $\rho$) between $B_{12}$ biosynthesis traits diversity and geo-environmental factors in whole layers (A) and in the SRF (B), DCM (C), MES (D) layers. Both taxonomic and functional trait diversity were analyzed. Geo-environmental factors including sampling depth, temperature, oxygen, nitrate, $NO_2NO_3$, and $PO_4$ concentrations were included.

**Supplementary Figure 8**. Associations between $B_{12}$ biosynthesis trait diversity indices and *metH* gene relative abundances in the global ocean.

**Supplementary Figure 9.** Random forest prediction of chlorophyll *a* concentrations using potential $B_{12}$ biosynthesis traits. Both the taxonomic (A) and functional trait (B) profiles were used for chlorophyll *a* concentration prediction. In the left panel, the SRF samples were used as the training dataset to predict chlorophyll *a* concentrations in DCM samples. In the right panel, the DCM samples were used as the training dataset to predict chlorophyll *a* concentrations in SRF samples.

### Supplementary Tables

**Supplementary Table 1.** Statistical testing of $B_{12}$ biosynthesis functional traits differences for samples in different ocean layers.

**Supplementary Table 2.** Microbial taxa potentially involved in $B_{12}$ biosynthesis in the global ocean.

**Supplementary Table 3.** Summary of microbial taxa carrying $B_{12}$ biosynthesis genes in ocean samples.

**Supplementary Table 4.** Summary of $B_{12}$ biosynthesis functional traits in ocean samples.

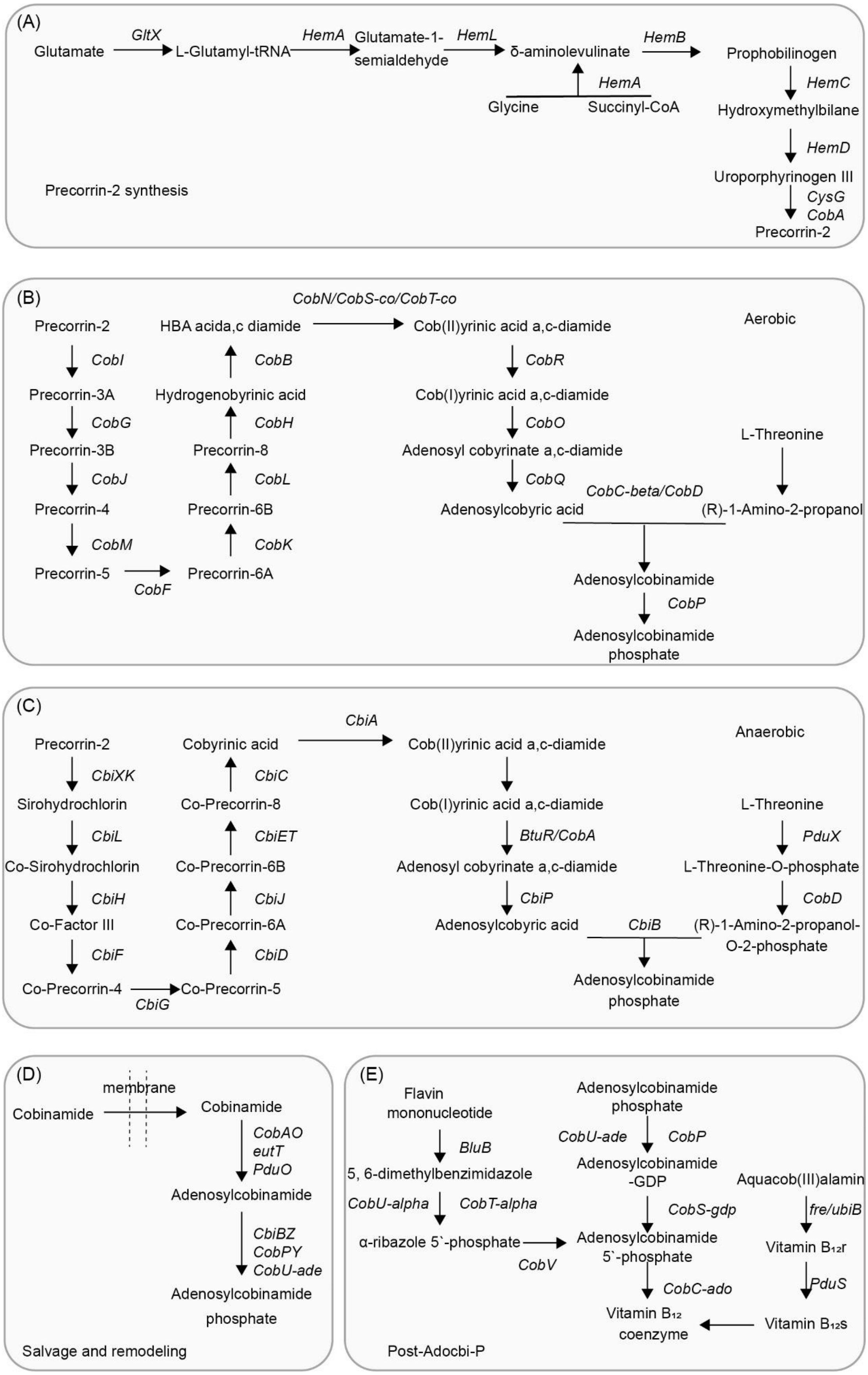


**Supplementary Figure 1.** Cobalamin biosynthesis pathway and related gene families. The $B_{12}$

biosynthesis pathway contains five modules, including precorrine-2 synthesis, aerobic pathway, anaerobic pathway, salvage and remodeling pathway, and post-AdoCbi-P pathway.

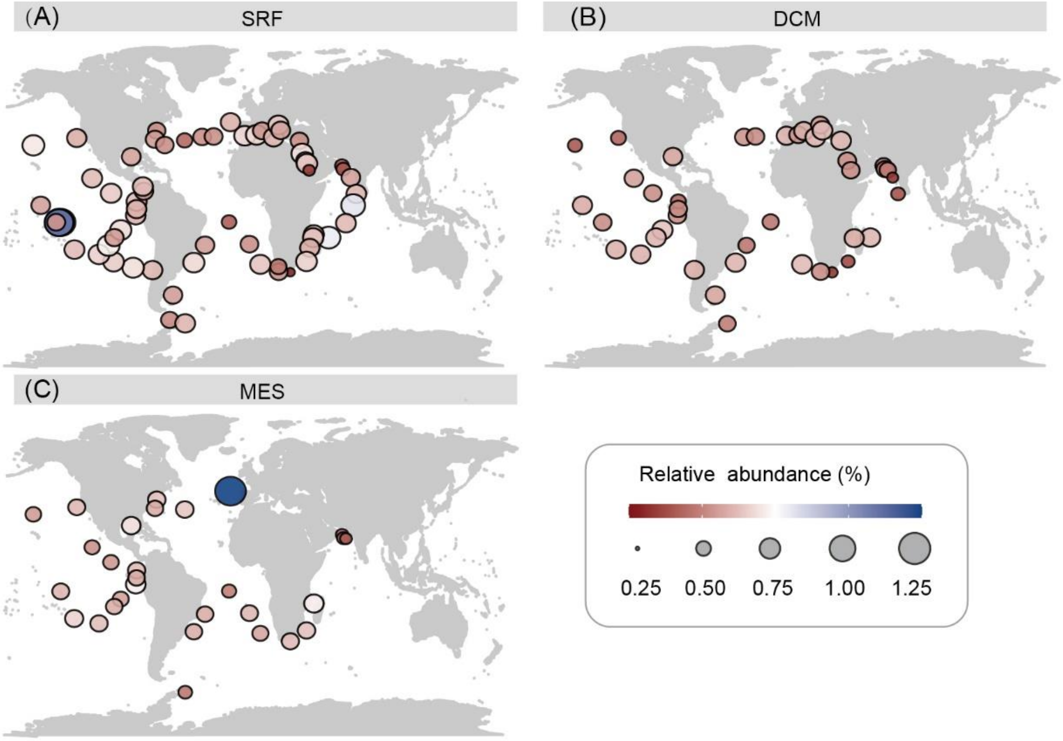


**Supplementary Figure 2.** Relative abundances of $B_{12}$ biosynthesis traits in the ocean. The size of the circles and colors represents the relative abundance of microbial functional genes involved in $B_{12}$ biosynthesis.

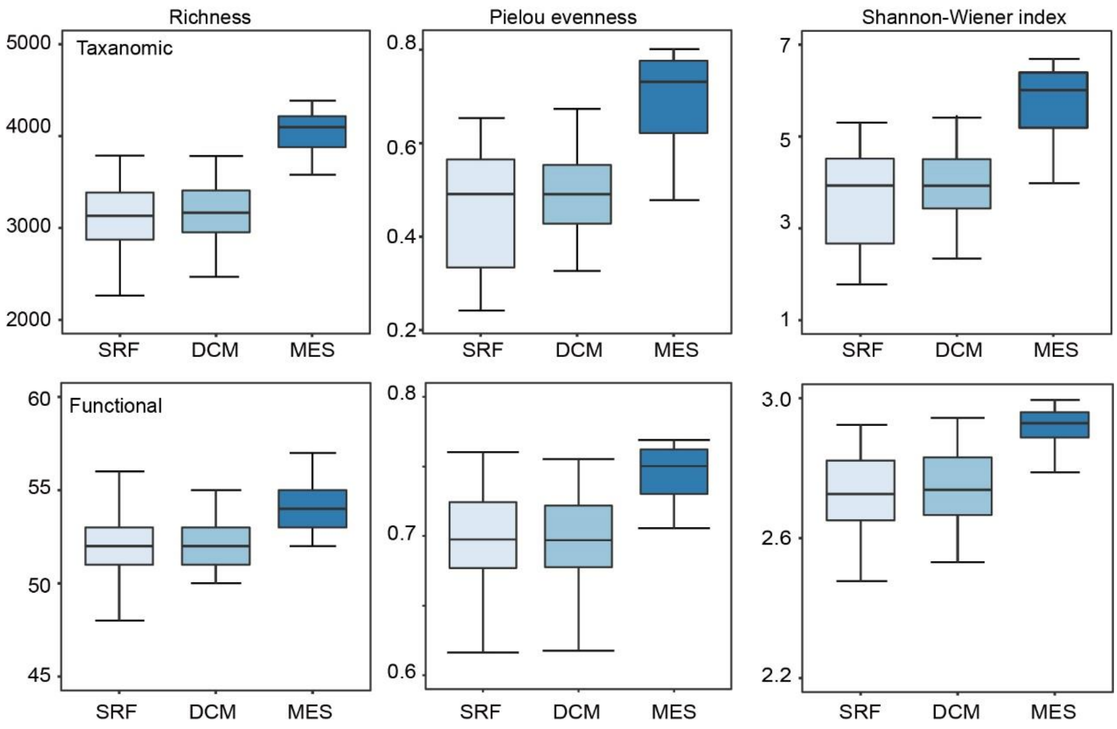


**Supplementary Figure 3.** Diversity indices (richness, evenness, and Shannon-Wiener index) of microbial communities carrying $B_{12}$ biosynthesis in SRF, DCM, and MES layers. Both the diversity indices for taxonomic groups (top) and functional traits (bottom) were presented.

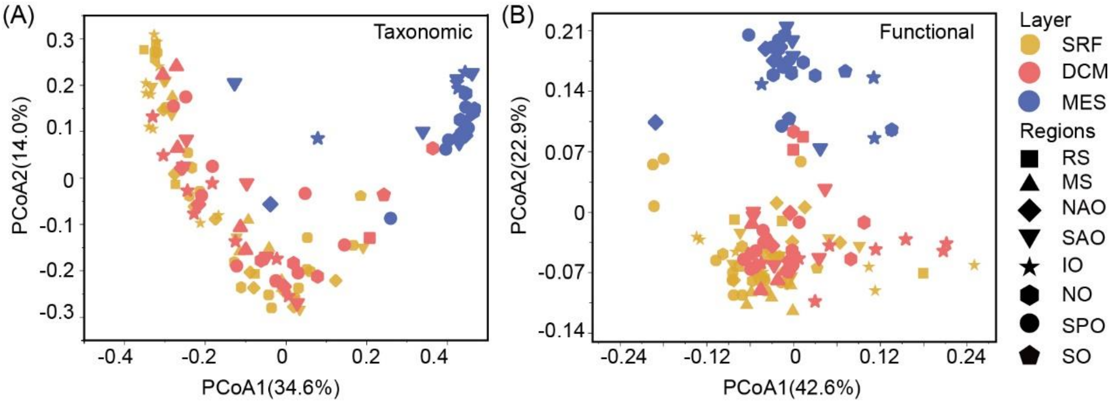


**Supplementary Figure 4.** PCoA clustering of taxonomic and functional trait potentially involved in $B_{12}$ biosynthesis in the global ocean. Bray-Curtis dissimilarity was used for distance calculation. Different colors represent samples in different ocean layers, while different shapes represent different oceans.

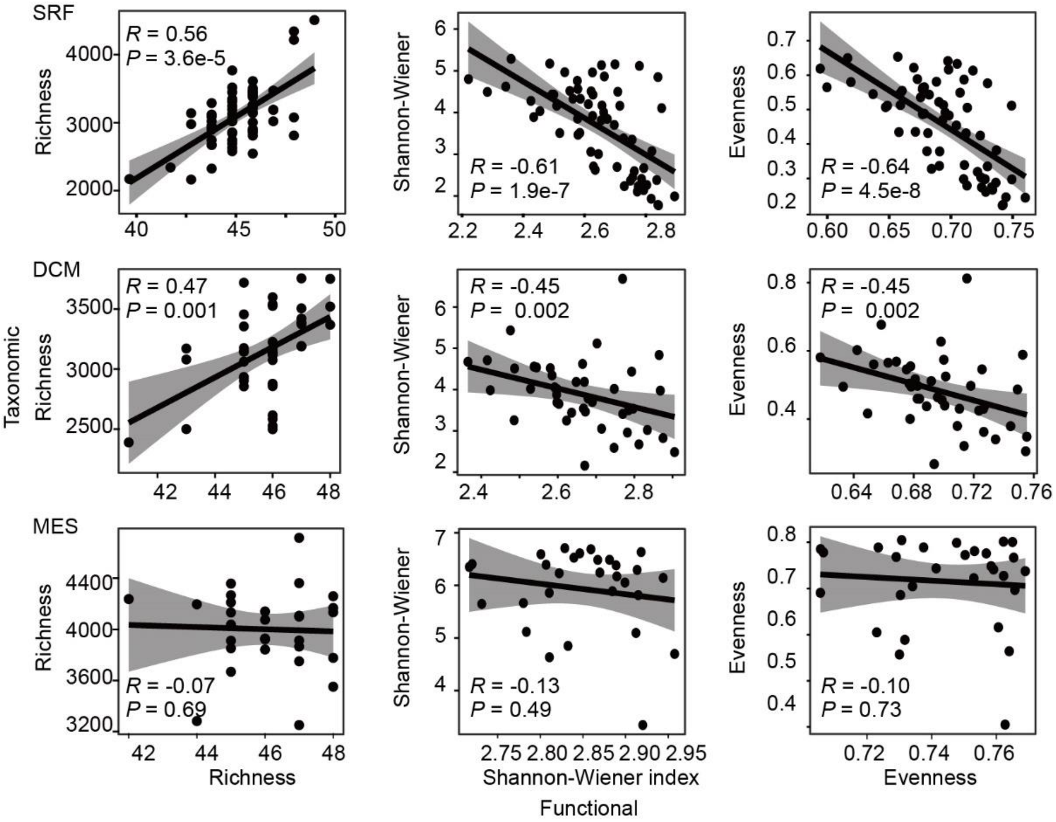


**Supplementary Figure 5.** Relationship between $B_{12}$ biosynthesis functional traits and their carrying taxa in different oceanic layers. Diversity indices including richness, evenness, and Shannon-Wiener index were analyzed.

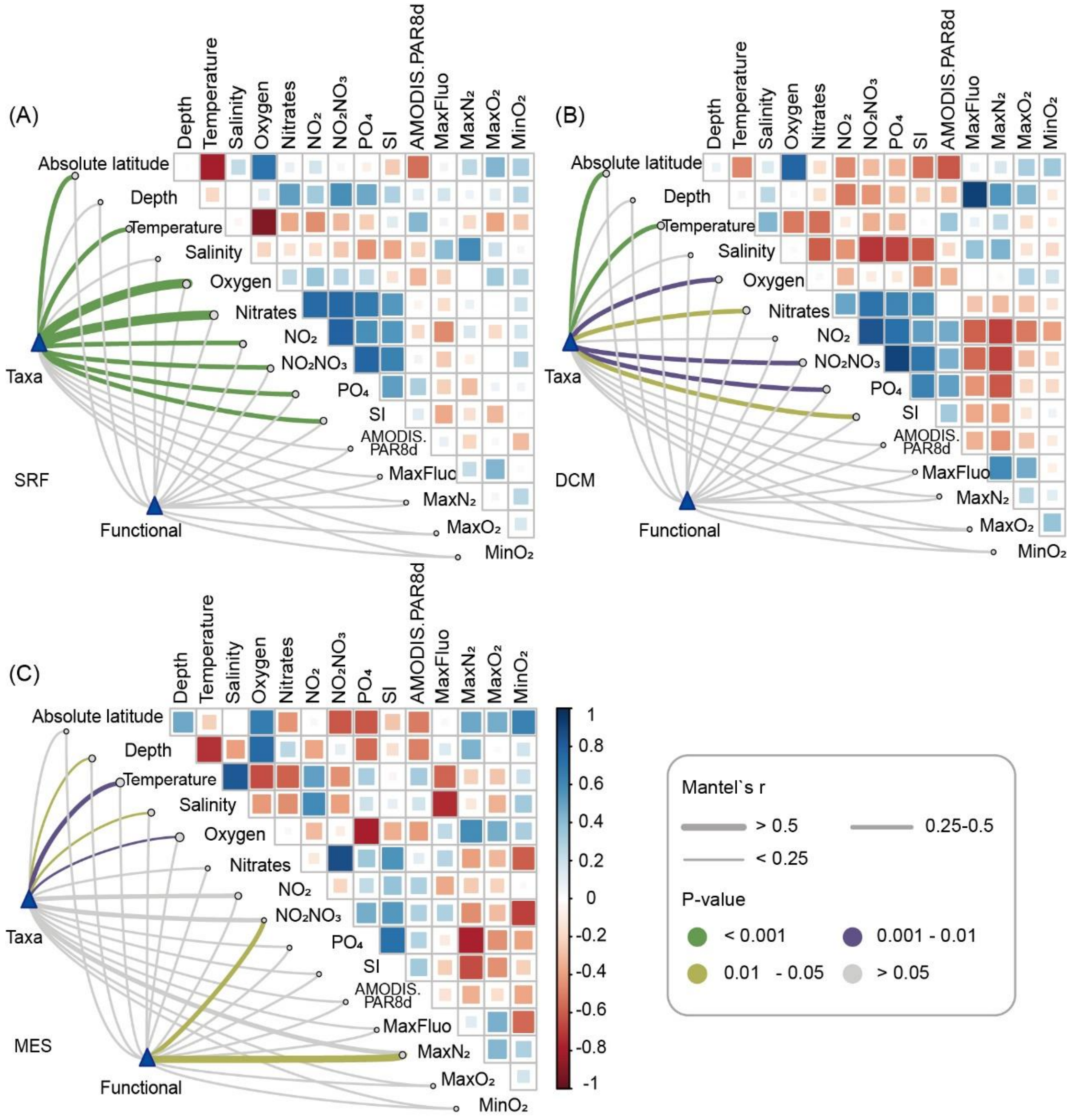


**Supplementary Figure 6.** Linking environmental factors with the compositional variations of $B_{12}$ biosynthesis traits. Pairwise comparisons of environmental factors are shown, with color gradients denoting Spearman's correlation coefficients. The $B_{12}$ biosynthesis traits in the SRF (A), DCM (B), and MES (C) layers were related to each environmental factor by partial (geographic distance–corrected) Mantel tests. Edge width corresponds to Mantel's r statistic, edge color corresponds to the statistical significance based on 9,999 permutations. Taxa: taxonomic composition; Fun: functional composition.

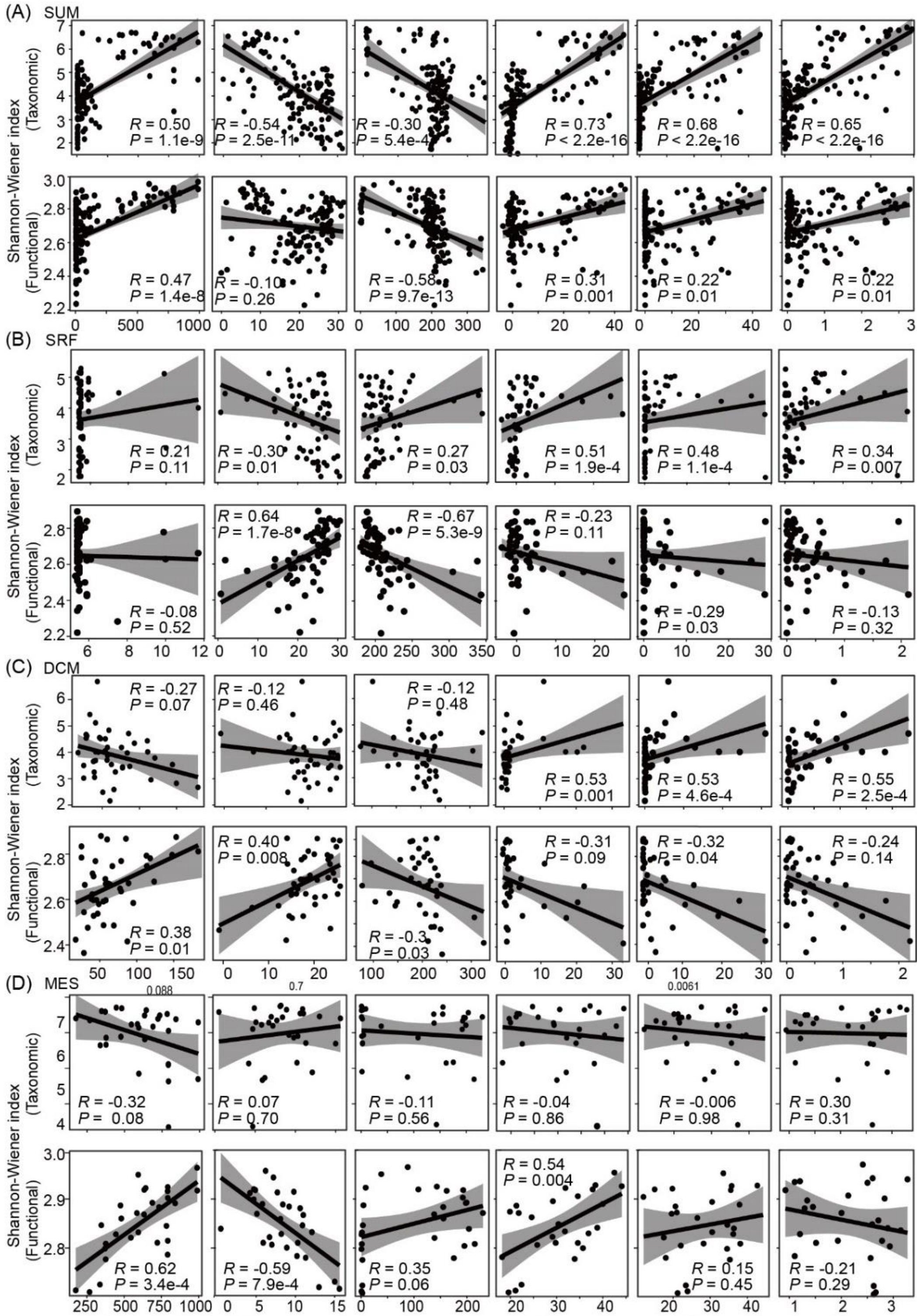


**Supplementary Figure 7.** Associations (Spearman's $\rho$) between $B_{12}$ biosynthesis traits diversity and geo-environmental factors in whole layers (A) and in the SRF (B), DCM (C), MES (D) layers. Both taxonomic and functional trait diversity were analyzed. Geo-environmental factors including sampling depth, temperature, oxygen, nitrate, $NO_2NO_3$, and $PO_4$ concentrations were included.

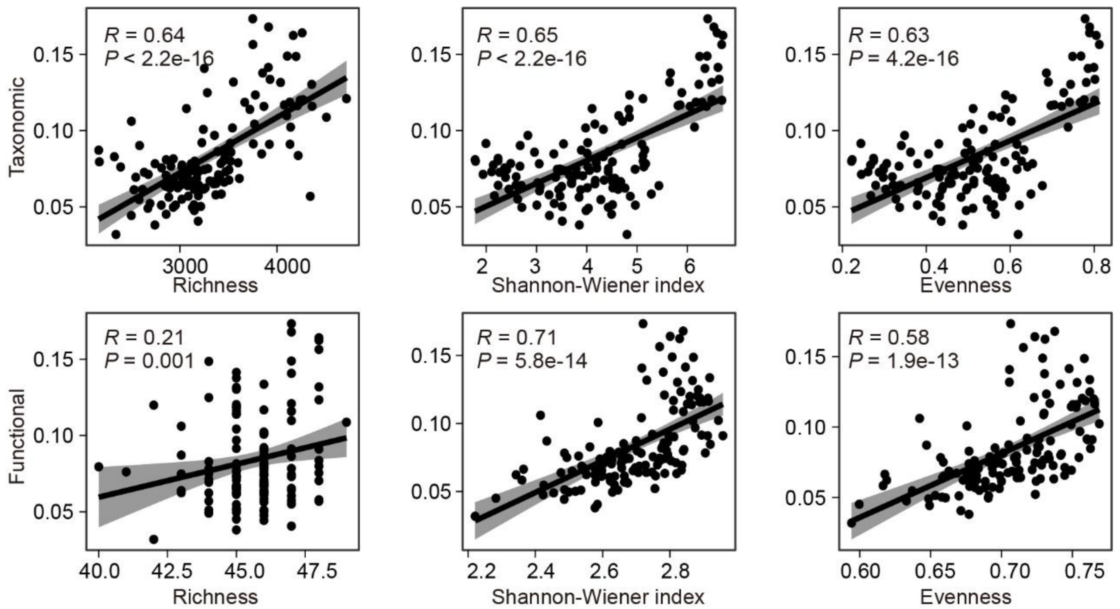


**Supplementary Figure 8.** Associations between $B_{12}$ biosynthesis trait diversity indices and *metH* gene relative abundances in the global ocean.

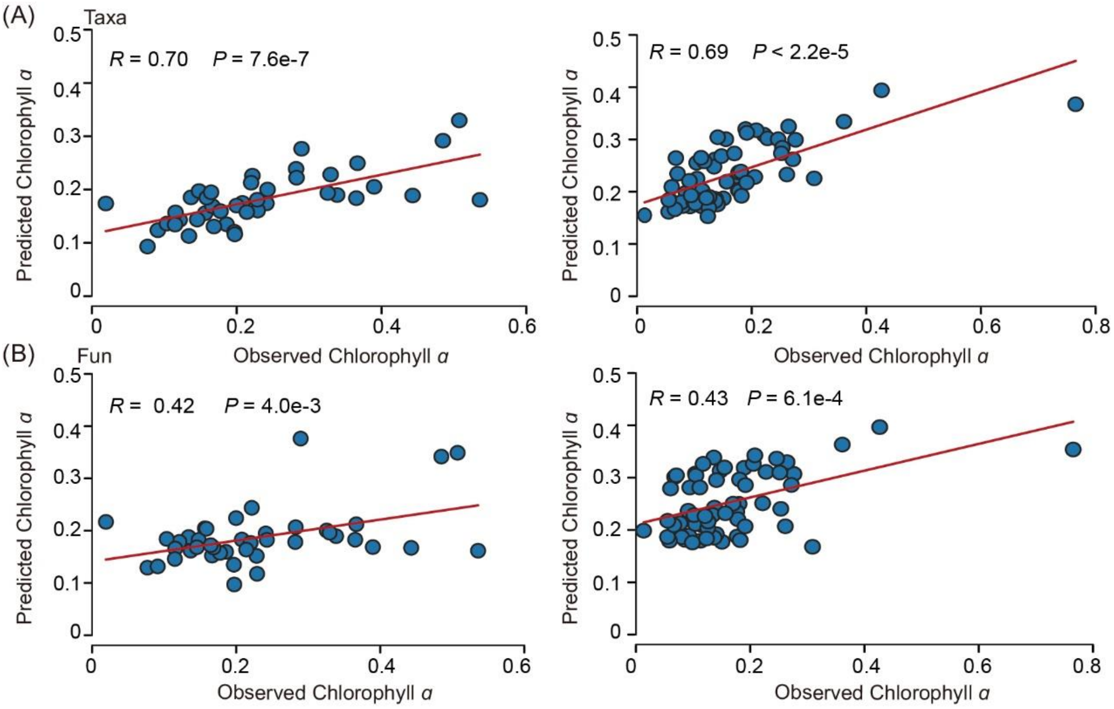


**Supplementary Figure 9.** Random forest prediction of chlorophyll *a* concentrations using potential $B_{12}$ biosynthesis traits. Both the taxonomic (A) and functional trait (B) profiles were used for chlorophyll *a* concentration prediction. In the left panel, the SRF samples were used as the training dataset to predict chlorophyll *a* concentrations in DCM samples. In the right panel, the DCM samples were used as the training dataset to predict chlorophyll *a* concentrations in SRF samples.

**Supplementary Table 1.** Statistical testing of $B_{12}$ biosynthesis functional traits differences for samples in different ocean layers. Three different permutation tests were performed, including the multiple response permutation procedure (MRPP), analysis of similarity (ANOSIM) and permutational multivariate analysis of variance (PERMANOVA). The Bray-Curtis dissimilarity was used for community dissimilarity. Red values indicate $p < 0.05$.

| | Taxonomic | | | | | | Functional | | | | | |
|---|---|---|---|---|---|---|---|---|---|---|---|---|
| | PERMANOVA | | ANOSIM | | MRPP | | PERMANOVA | | ANOSIM | | MRPP | |
| | F | *P* | R | *P* | Delta | *P* | F | *P* | R | *P* | Delta | *P* |
| SRF vs. DCM | 1.485 | 0.050 | 0.007 | 0.345 | 0.217 | 0.038 | 1.689 | 0.185 | -0.015 | 0.709 | 0.027 | 0.102 |
| SRF vs. MES | 28.866 | 0.001 | 0.627 | 0.001 | 0.205 | 0.001 | 13.406 | 0.001 | 0.123 | 0.013 | 0.027 | 0.001 |
| DCM vs. MES | 24.272 | 0.001 | 0.674 | 0.001 | 0.198 | 0.001 | 8.560 | 0.001 | 0.139 | 0.001 | 0.026 | 0.001 |
